# The Role of Central Amygdala Corticotropin-Releasing Factor in Predator Odor Stress-Induced Avoidance Behavior and Escalated Alcohol Drinking in Rats

**DOI:** 10.1101/870386

**Authors:** Marcus M. Weera, Allyson L. Schreiber, Elizabeth M. Avegno, Nicholas W. Gilpin

**Author notes:** Equal contributions (Marcus M. Weera, Allyson L. Schreiber). Corresponding author: Marcus M. Weera, Ph.D., Department of Physiology, 1901 Perdido Street, New Orleans, LA 70112.

## Abstract

Post-traumatic stress disorder (PTSD) is characterized by avoidance of trauma-associated stimuli and amygdala hyperreactivity, and is highly co-morbid with alcohol use disorder (AUD). Our lab uses a predator odor (bobcat urine) stress model that produces conditioned avoidance of an odor-paired context in a subset of rats, mirroring avoidance symptoms that manifest in some but not all humans exposed to trauma. We previously showed that after predator odor stress, Avoiders exhibit escalated alcohol drinking, higher aversion-resistant operant alcohol responding, hyperalgesia, and greater anxiety-like behavior compared to unstressed Controls. We also showed that systemic antagonism of corticotropin-releasing factor-1 receptors (CRFR1) reduced escalation of alcohol drinking in rats not indexed for avoidance, that corticotropin-releasing factor (CRF) infusions into the central amygdala (CeA) produced conditioned place avoidance in stress-naïve rats, and that intra-CeA infusion of a CRFR1 antagonist reduced hyperalgesia in Avoiders. Here, we show that avoidance behavior is persistent after repeated predator odor exposure and is resistant to extinction. In addition, Avoiders showed lower weight gain than Controls after predator odor re-exposure. In the brain, higher avoidance was correlated with higher number of c-Fos+ cells and CRF immunoreactivity in the CeA. Finally, we show that intra-CeA CRFR1 antagonism reversed post-stress escalation of alcohol drinking and reduced avoidance behavior in Avoiders. Collectively, these findings suggest that elucidation of the mechanisms by which CRFR1-gated CeA circuits regulate avoidance behavior and alcohol drinking may lead to better understanding of the neural mechanisms underlying co-morbid PTSD and AUD.

## INTRODUCTION

Post-traumatic stress disorder (PTSD) has a prevalence rate of ~8% in the general population and ~20% among U.S. combat veterans [1,2] and costs the U.S. billions of dollars annually [3]. According to the Diagnostic and Statistical Manual of Mental Disorders (5^th^ edition), a PTSD diagnosis requires presence of symptoms from all four of the following symptom clusters for at least one month: 1) intrusive recollections of the traumatic event, 2) avoidance of trauma-related stimuli, 3) negative alterations in cognition, and 4) alterations in arousal and mood. Of these symptom clusters, avoidance of trauma-related stimuli may be the most detrimental to psychosocial functioning [4,5], is a strong predictor of future PTSD severity [6–8] and poor treatment response [9], and is predicted by amygdala hyperreactivity [10]. Critically, PTSD is highly co-morbid with alcohol use disorder (AUD), with 22-43% of individuals with PTSD meeting criteria for AUD, compared to 8% in the general population [11]. Relative to either disorder alone, co-morbid AUD and PTSD produce worse outcomes, such as increased incidence of disease and greater loss of work productivity [12].

The use of animal models to investigate the neurobiological factors underlying traumatic stress effects is critical for the development of better prevention, diagnostic, and treatment strategies. Our lab uses a model in which a subset of rats exposed to predator odor (bobcat urine) show persistent avoidance (up to 6 weeks) of odor-paired stimuli (context with distinct tactile and visual cues), mirroring avoidance symptoms that manifest in some but not all humans exposed to traumatic stress [13,14]. After predator odor stress exposure, rats classified as Avoiders exhibit escalated alcohol drinking, more aversion-resistant operant alcohol responding [15], hyperalgesia [16], and higher anxiety-like behavior (also seen in Non-Avoiders) than unstressed Controls [17]. Therefore, this animal model may be useful for understanding the neurobiology underlying behavioral changes after traumatic stress.

One goal of this study was to further characterize the avoidance phenotype in our predator odor stress model. In other predator stress models, rats exposed to a worn cat collar exhibit persistent and extinction-resistant reduction in locomotor activity after repeated exposure to the predator odor stress [18]. In addition, repeated cat exposures reduce weight gain and produce adrenal hypertrophy in rats [19]. Here, we tested avoidance behavior after repeated exposures to predator odor, the effects of repeated exposure to predator odor-paired context (i.e., extinction sessions) on avoidance behavior, and the effect of predator odor exposure on body weight gain.

Another goal of this study was to elucidate the neurobiological mechanisms contributing to predator odor-paired context avoidance and post-stress alcohol drinking in the bobcat urine stress model. We previously found that intra-CeA corticotropin releasing factor (CRF) infusions produce dose-dependent avoidance of the CRF-paired chamber in stress-naïve rats, intra-CeA CRF-1 receptor (CRFR1) antagonism reduces hyperalgesia in Avoiders after stress [16], and systemic CRFR1 antagonism reduces escalated alcohol drinking in rats not indexed for avoidance [20]. Here, we tested the hypotheses that Avoiders would exhibit greater CeA activation (as indicated by higher number of c-Fos-positive cells) and higher CeA CRF immunoreactivity (CRF-ir) than Non-Avoiders and unstressed Controls. We also tested the hypotheses that CRFR1 blockade would reduce avoidance of a predator odor-paired context and post-stress alcohol drinking in Avoiders.

## METHODS

All procedures were approved by the Institutional Animal Care and Use Committee of the Louisiana State University Health Sciences Center and were in accordance with the National Institute of Health guidelines. General methods are detailed in the supplementary material.

### Predator odor place conditioning

Rats underwent a 5-day predator odor conditioned place avoidance (CPA) procedure as previously described [21]. Briefly, on the first day, rats were allowed access to three chambers with distinct tactile (circles vs. grid. vs. rod floor) and visual (circles vs. white vs. stripes) cues (3-chamber Pretest). For each rat, the chamber with the most deviant time relative to the other two chambers (i.e., most extremely preferred or avoided chamber) was excluded from all subsequent testing. On day 2, rats were allowed 5 min to explore the two non-excluded chambers (2-chamber Pretest). Rats were assigned to predator odor stress or unstressed control groups that were counterbalanced for magnitude of baseline preference across the two chambers being used for each animal. For rats in the stress group, an unbiased and counterbalanced procedure was used to determine which chamber (i.e., more preferred or less preferred) would be paired with predator odor exposure in individual rats. On day 3, rats were placed in one chamber without odor for 15 min (neutral conditioning). On day 4, rats were placed in the other chamber for 15 min in the presence (or absence for unstressed Controls) of a bobcat urine-soaked sponge placed under the floor of the chamber (predator odor conditioning). On day 5, rats were again allowed to explore the two chambers for 5 min (Posttest). Time spent in each chamber was scored by an observer. Rats that showed >10 s decrease in time spent in the odor-paired context between 2-chamber pretest and posttest were classified as ‘Avoiders,’ and all other odor-exposed rats were classified as ‘Non-Avoiders’. In some experiments, there was more than one Posttest (see below).

### Repeated predator odor exposure effect on avoidance behavior

Rats underwent the predator odor CPA procedure and were indexed for avoidance (Controls, n=6; Non-Avoiders, n=9; Avoiders, n=6). Rats were re-exposed to the neutral chamber (5 days after Posttest) and to predator odor in the CS+ chamber (or no odor for Controls; 6 days after Posttest) and re-indexed for avoidance the next day (**Fig. 1A**).

**Figure 1.**
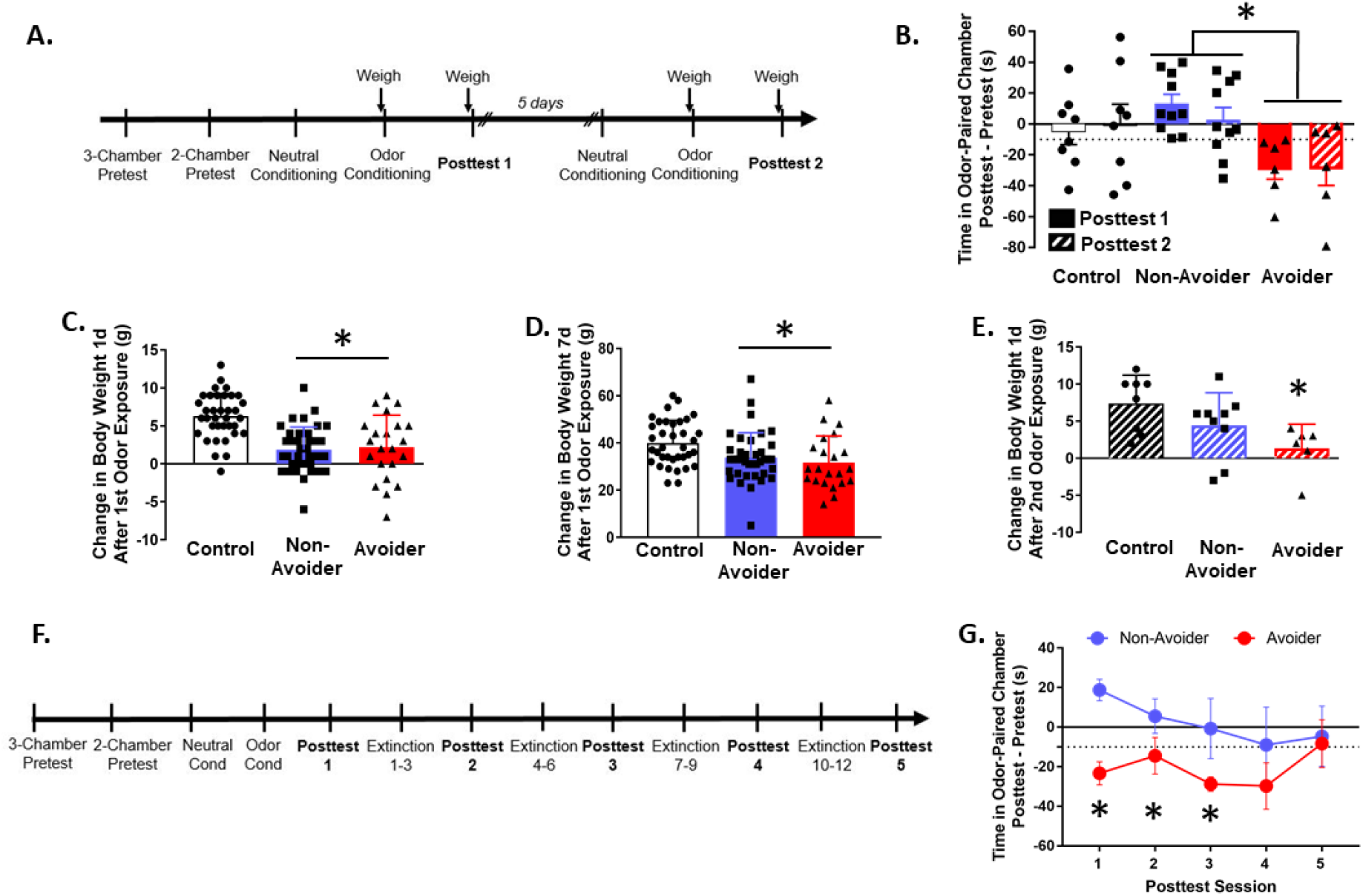
Predator odor stress produces avoidance behavior that is stable and resistant to extinction and changes in weight gain. **(A)** Timeline for testing the effect of repeated predator odor exposure on avoidance behavior and weight gain. **(B)** Time in odor-paired chamber during Posttests 1 (solid bars) and 2 (hatched bars) (after 1^st^ and 2^nd^ predator odor exposure). \**p* = 0.01, Avoiders vs. Non-Avoiders. **(C)** Change in body weight 1 day after predator odor exposure and **(D)**7 days after predator odor exposure. \**p* ≤ 0.01, stressed rats (Avoiders and Non-Avoiders combined) vs. Controls. **(E)** Change in body weight 1 day after 2^nd^ predator odor exposure. \**p* = 0.04, Avoiders vs. Controls. **(F)** Timeline for testing the effect of repeated extinction sessions on avoidance behavior. **(G)** Time in odor-paired chamber during Posttests 1-5. Each post-test was separated by 3 extinction sessions on consecutive days between post-tests. \**p*’s < 0.05, Avoiders vs. Non-Avoiders.

### Predator odor exposure effect on body weight gain

Rats underwent the predator odor CPA procedure and were indexed for avoidance (Controls, n=36; Non-Avoiders, n=38; Avoiders, n=22). A subset of rats (Controls, n=7; Non-Avoiders, n=9; Avoiders, n=6) was re-exposed to predator odor (or no odor for unstressed Controls) in the CS+ chamber (or no odor for Controls) 7 days after the 1^st^ odor exposure (6 days after Posttest) (**Fig. 1A**). Rats were weighed immediately before and 1 day after the first predator odor exposure, then again immediately before and 1 day after the 2^nd^ predator exposure. We calculated change in body weight 1 day and 7 days after the 1^st^ odor exposure, as well as 1 day after the 2^nd^ odor exposure. A subset of rats were sacrificed 7 days after the 1st odor exposure and their adrenal and thymus glands were weighed.

### Extinction of avoidance of predator odor-paired context

Rats underwent the predator odor CPA procedure and were indexed for avoidance (Non-Avoiders, n=6; Avoiders, n=4). After posttest 1, rats were placed in the neutral chamber for 15 min in the morning and in the odor-paired chamber (in the absence of odor) for 15 min in the afternoon over the next 3 consecutive days (i.e., 3 extinction sessions). On the day after the 3^rd^ extinction session, rats were re-indexed for avoidance (posttest 2). This entire procedure was repeated 3 more times, such that each rat underwent 5 total posttests, each of them (except the first one) preceded by 3 extinction days (**Fig. 1F**).

### Predator odor exposure effect on c-Fos and CRF immunoreactivity in CeA

Rats were subjected to the predator odor CPA procedure and indexed for avoidance (Controls, n=12; Non-Avoiders, n=13; Avoiders, n=6). Six days after posttest, all Non-Avoiders and Avoiders were re-exposed to predator odor for 15 min in the CS+ chamber; half of the unstressed Controls (n=6) were also stressed that day (denoted as the “Single Stress” group) whereas the other half of unstressed Controls (n=6) were not exposed to odor and remained stress-naïve. Rats were sacrificed 90 min after 2^nd^ odor exposure and brains were processed for immunohistochemical labeling for c-Fos and CRF.

### Intra-CeA CRFR1 antagonism effect on post-stress avoidance and alcohol drinking

Please refer to Figure 4A for experimental timeline. Rats were trained to orally self-administer 10% (w/v) ethanol or water as previously described [21]. Upon stabilization of operant responding in 30 min sessions, rats were implanted with bilateral cannulae targeting the CeA. After ~1 week of recovery, rats were allowed to resume alcohol self-administration. Upon stabilization of responding, self-administration following intra-CeA vehicle infusion was recorded. Subsequently, rats underwent the predator odor CPA procedure and were indexed for avoidance (Control, n=14; Non-Avoider, n=27; Avoider, n=19). On days 2, 5, 8, 10, and 12 after posttest, rats were allowed to self-administer alcohol in 30-min operant sessions that were immediately preceded by intra-CeA infusion of a CRFR1 antagonist, R121919 (0.25 μg/0.5 μL/side), or vehicle (20% w/v 2-hydroxypropyl cyclodextrin in saline). One day after the last session of alcohol self-administration, rats were re-indexed for avoidance (Posttest 2) to test the effect of repeated intra-CeA R121919 or vehicle infusions on avoidance behavior. Two days later, a subset of rats (Control, n=6; Non-Avoiders, n=11; Avoiders, n=12) was re-indexed for avoidance (posttest 3) immediately following intra-CeA infusion of R121919 or vehicle to test the acute effect of CRFR1 antagonism on avoidance behavior. Rats with infusions outside the CeA were removed from all analyses.

### Statistical Analysis

Data in Figures 1–4 are shown as mean ± SEM. All statistical analyses were performed using SPSS 25 (IBM Corp., Armonk, NY). Data were analyzed using two-way, one-way, or repeated measures ANOVAs, and significant effects were followed-up with Fisher’s LSD post-hoc tests. The specific statistical tests used for each experiment are reported in the Results. Outliers were detected using the 1.5 x IQR rule and animals with outlier values were excluded from all behavioral and molecular data analysis (n=12 across all experiments). Statistical significance was set at *p* < 0.05.

## RESULTS

### Avoidance behavior is similar after 1^st^ and 2^nd^ predator odor exposure

Rats were exposed to predator odor twice and indexed for avoidance after each exposure (**Fig. 1A**). Time spent in the odor-paired chamber during Posttest 1 (after first odor exposure) and Posttest 2 (after 2^nd^ odor exposure) were similar for Avoiders and Non-Avoiders (repeated measures Posttest x Stress Group ANOVA, *p* = 0.76). Across both Posttest sessions, time spent in odor-paired chamber was significantly different between Avoiders and Non-Avoiders (main effect of Stress Group, *F*_2,21_ = 4.93, *p* = 0.02; post-hoc test, *p* = 0.01) (**Fig. 1B**).

### Predator odor exposure reduces body weight gain

Both Avoiders and Non-Avoiders gained less weight than Controls 1 day (one-way ANOVA, *F*_2,93_ = 19.1, *p* < 0.001; post-hoc tests, *p*’s < 0.001; **Fig. 1C**) and 7 days (one-way ANOVA, *F*_2,93_ = 5.3, *p* = 0.01; post-hoc tests, *p*’s < 0.05; **Fig. 1D**) after the 1^st^ predator odor exposure. One day after the 2^nd^ predator odor exposure, only Avoiders showed reduced body weight gain relative to unstressed Controls (one-way ANOVA, effect of Stress Group, *F*_2,20_ = 4.1, *p* = 0.04; post-hoc test, *p* = 0.03; **Fig. 1E**). A subset of rats were sacrificed 7 days after odor exposure (**Fig. 1A**) and their adrenal and thymus glands were weighed. There were no differences in adrenal or thymus gland weights between Controls, Non-Avoiders, and Avoiders (data not shown).

### Avoidance behavior is resistant to extinction

Avoider and Non-Avoider rats were exposed repeatedly to the neutral and odor-paired chambers to test the persistence of avoidance behavior after extinction sessions (**Fig. 1F**). Time spent in the odor-paired chamber was significantly different between Avoiders and Non-Avoiders during Posttests 1-3 (t-tests, *p*’s < 0.05), but not during Posttests 4 and 5 (**Fig. 1G**).

### Avoiders have more c-Fos+ cells in CeA after odor re-exposure

Rats were exposed to predator odor, indexed for avoidance, re-exposed to predator odor, and sacrificed 90 min later for c-Fos immunostaining (**Fig. 2A**). Controls were never exposed to predator odor and Single Stress rats were exposed to predator odor once 90 min before sacrifice (see Methods). Avoiders had a higher number of c-Fos+ cells in the CeA than Controls, Non-Avoiders, and rats that received Single Stress exposure (one-way ANOVA, main effect of Stress Group, *F*_1,25_ = 3.1, *p* = 0.046; post-hoc tests, *p* = 0.030, 0.012, and 0.013, respectively; **Fig. 2B**).

**Figure 2.**
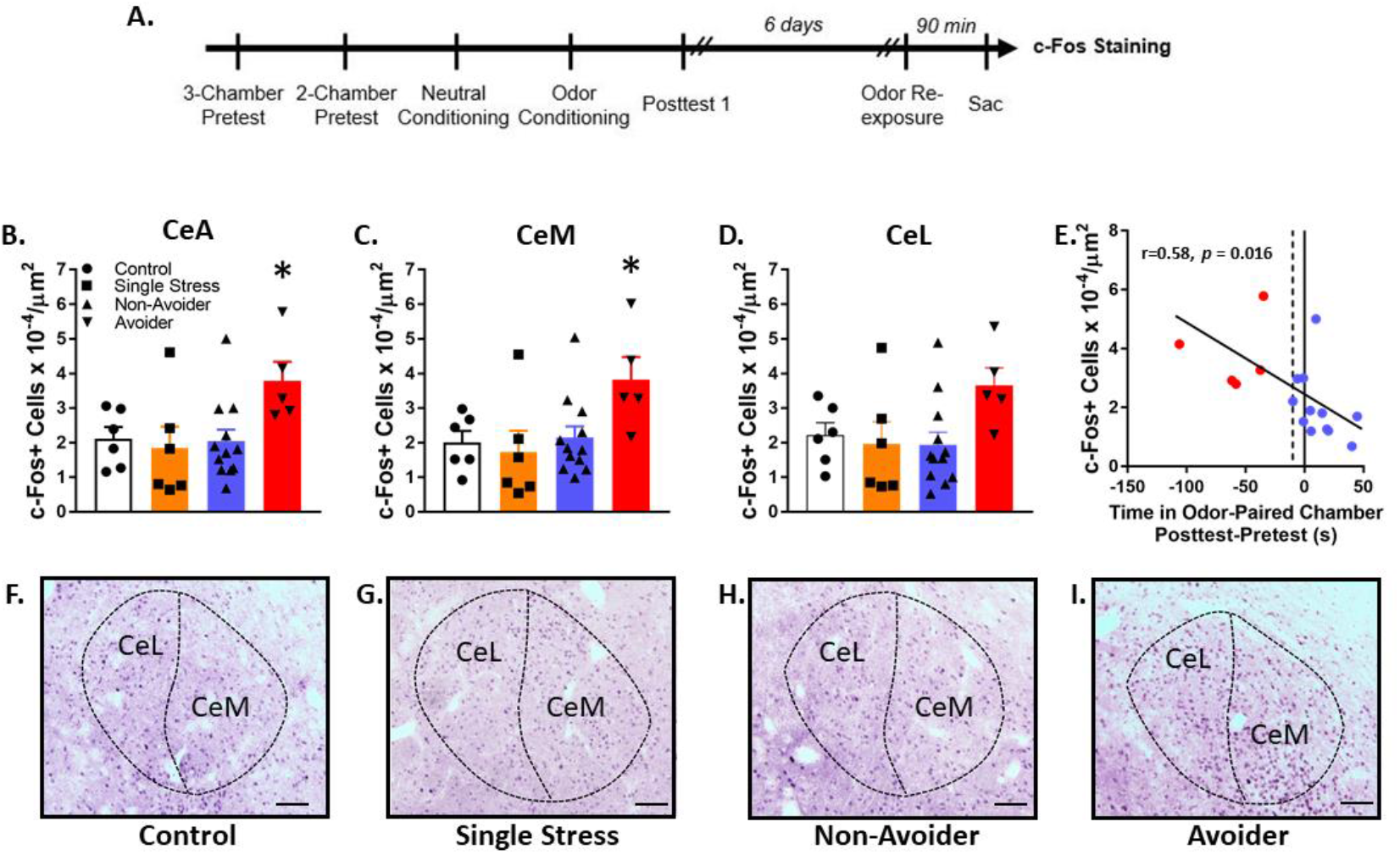
Avoiders have more c-Fos+ cells in CeA than Non-Avoiders. **(A)** Timeline for testing the effect of predator odor exposure on c-Fos induction. **(B)** c-Fos+ cells in CeA, **(C)** CeM, and **(D)** CeL. \**p*’s < 0.05, Avoiders vs. Non-Avoiders, Single Stress and Unstressed Controls. **(E)** Correlation between avoidance and number of c-Fos+ cells in CeA. **(F)** Representative images of c-Fos staining in Control, **(G)** Single Stress, **(H)** Non-Avoider, and **(I)** Avoider rats. Scale bar = 100 μm.

Analyses split by medial (CeM) and lateral (CeL) CeA subdivisions showed that in the CeM, c-Fos+ cell counts were higher in Avoiders than in Controls, Non-Avoiders, and rats that received Single Stress exposure (one-way ANOVA, main effect of Stress Group, *F*_1,25_ = 3.2, *p* = 0.039; post-hoc tests, *p* = 0.021, 0.016, and 0.009, respectively; **Fig. 2C**). However, in the CeL, the effect of Stress group was not significant (one-way ANOVA, *p* = 0.089; **Fig. 2D**). Across Avoider and Non-Avoider groups (i.e., all rats exposed to odor twice), there was a significant inverse correlation between number of c-Fos+ cells in CeA and time spent in the odor-paired chamber during Posttest (Pearson’s r = −0.58, *p* = 0.016; **Fig. 2E**), such that high avoidance was correlated with high number of c-Fos+ cells in CeA. Representative images of c-Fos staining are shown in **Fig. 2F-I**.

### Avoiders have higher CRF immunoreactivity in CeA after odor re-exposure

Rats were exposed to predator odor, indexed for avoidance, re-exposed to predator odor, and sacrificed 90 min later for CRF immunostaining (**Fig. 3A**). Avoiders showed greater CRF-ir in CeA than Non-Avoiders, Controls, and rats that received Single Stress exposure (one-way ANOVA, main effect of Stress Group, *F*_1,23_ = 4.5, *p* = 0.01; post-hoc tests, *p* = 0.002, 0.022, and 0.011, respectively; **Fig. 3B**). Analyses of CRF immunoreactivity in CeM and CeL individually did not yield statistically significant effects (**Fig. 3C, D**). In Avoiders and Non-Avoiders, there was a significant inverse correlation between CeA CRF immunoreactivity and time spent on the odor-paired chamber during Posttest (Pearson’s r = −0.67, *p* = 0.004; **Fig. 3E**), such that high avoidance was correlated with high CRF-immunoreactivity in CeA. Representative images of CRF staining are shown in **Fig. 3F-I**.

**Figure 3.**
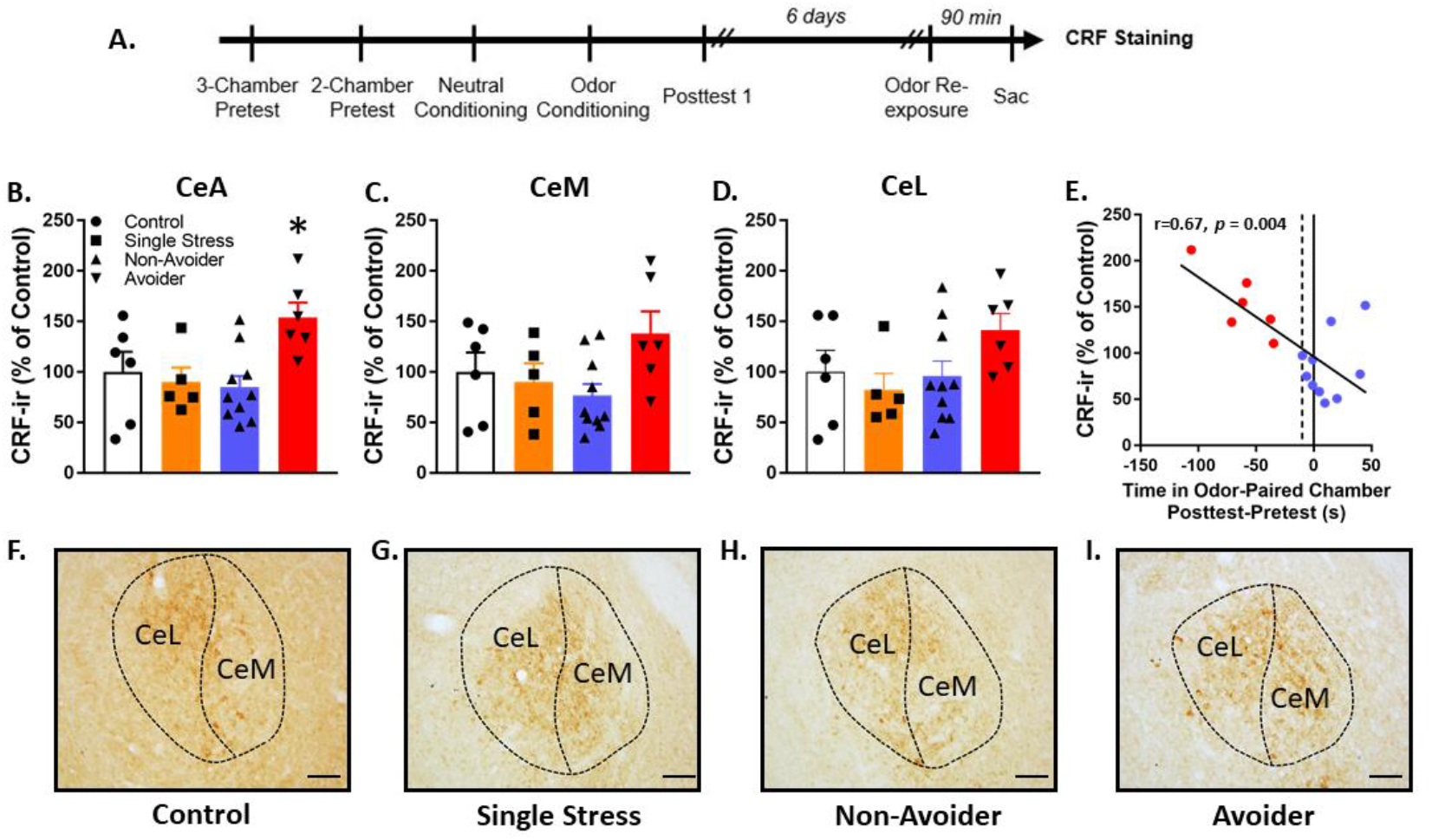
Avoiders have greater CRF immunoreactivity in CeA than Non-Avoiders. **(A)** Timeline for testing the effect of predator odor exposure on CRF immunoreactivity. **(B)** CRF immunoreactivity in CeA, **(C)** CeM, and **(D)** CeL. \**p*’s < 0.05, Avoiders vs. Non-Avoiders, Single Stress and Controls. **(E)** Correlation between avoidance and CeA CRF immunoreactivity. **(F)** Representative images of CRF staining in CeA of Control, **(G)** Single Stress, **(H)** Non-Avoider, and **(I)** Avoider rats. Scale bar = 100 μm.

**Figure 4.**
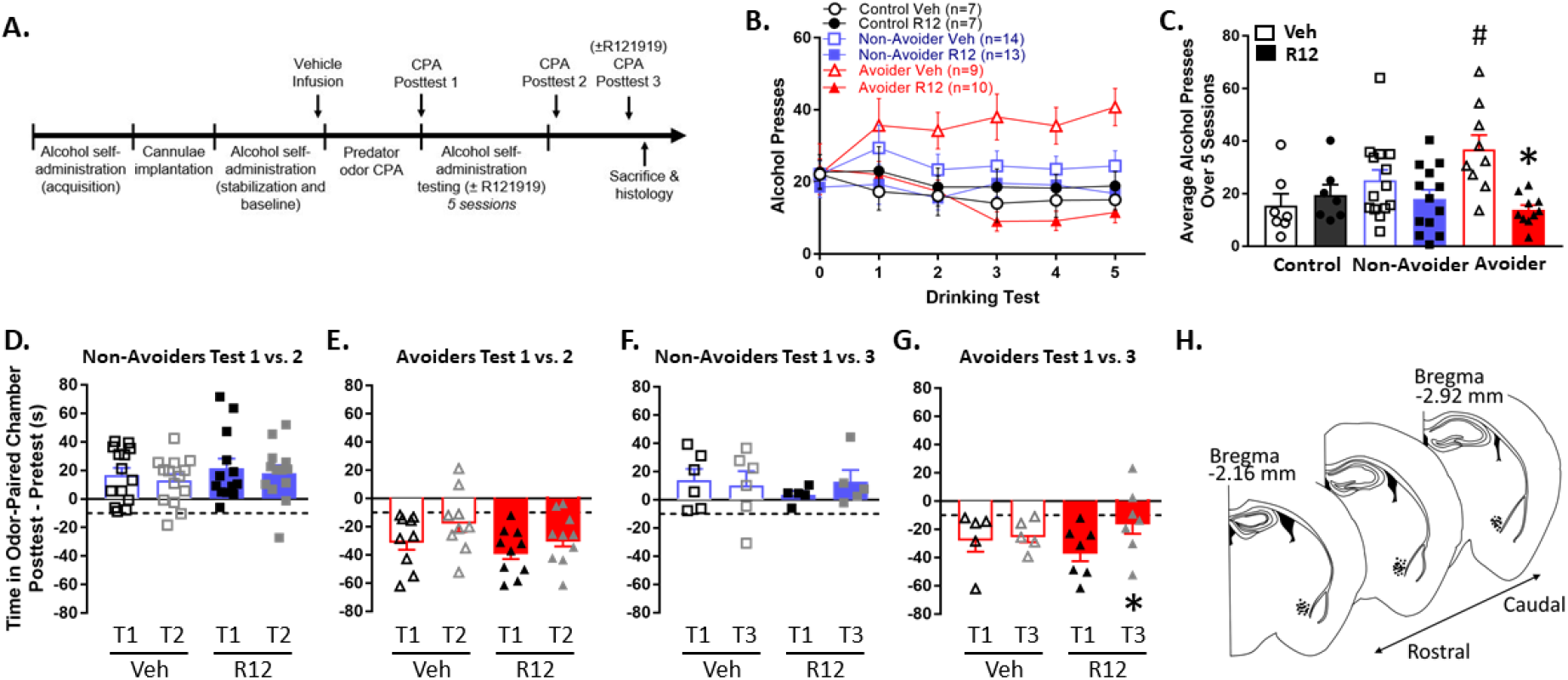
Intra-CeA CRFR1 antagonism reduces escalated alcohol drinking in Avoiders and avoidance behavior. **(A)** Timeline for testing the effect of intra-CeA CRFR1 antagonism on alcohol drinking and avoidance behavior. **(B)** Number of alcohol presses across 5 x 30 min operant alcohol oral self-administration test sessions. Rats were treated with R121919 (0.25 μg/side) or vehicle immediately before test sessions 1-5. All rats were treated with vehicle immediately before test session 0. **(C)** Number of alcohol presses averaged across test sessions 1-5. ^#^*p* = 0.008, Avoiders vs. Controls in the Vehicle group. \**p* = 0.001, R121919 vs. Vehicle treatment in the Avoider group. **(D)** Time spent in odor-paired chamber during Posttests 1 and 2 by Non-Avoiders and **(E)** Avoiders. All rats received intra-CeA vehicle infusions 1 day prior to Posttest 1 and were treated repeatedly with intra-CeA R121919 or vehicle during the 5 intervening operant alcohol self-administration sessions between Posttests 1 and 2 (last infusion occurred 1 day prior to Posttest 2). **(F)** Time spent in odor-paired chamber during Posttests 1 and 3 by Non-Avoiders and **(G)** Avoiders. Rats were treated with intra-CeA R121919 or vehicle immediately before Posttest 3. \**p* = 0.003, Avoiders treated with R121919, Posttest 1 vs. 3. **(H)** Schematic of infusion sites within the CeA.

### Intra-CeA CRFR1 antagonism reduces escalated alcohol drinking in Avoiders

Rats were trained to orally self-administer alcohol, exposed to predator odor, indexed for avoidance, and allowed to self-administer alcohol during five 30-min sessions on days 2, 5, 8, 10, and 12 after Posttest (**Fig. 4A**). Rats were pre-treated with intra-CeA infusions of R121919 (CRFR1 antagonist; 0.25 μg/side) or equivalent volume of vehicle immediately before operant self-administration testing sessions. A comparison of the average number of alcohol presses over 5 testing sessions between Avoiders and Controls in the Vehicle group showed significant stress-induced escalation of alcohol intake in Avoiders (two-way ANOVA, Stress Group x Treatment interaction, *F*_*2,54*_ = 4.7, *p* = 0.013; post-hoc test, *p* = 0.008). Within the Avoider group, R121919 treatment reduced the number of alcohol presses across 5 testing sessions relative to Vehicle-treated animals (effect of Treatment, *F*_1,17_ = 17.4, *p* = 0.001; **Fig. 4B, C**).

### Intra-CeA CRFR1 antagonism reduces avoidance of the predator odor-paired context

Rats were re-indexed for avoidance behavior 1 day after the last alcohol self-administration testing session in the absence of drug treatment (Posttest 2). Relative to Posttest 1, neither Avoiders nor Non-Avoiders showed changes in time spent in odor-paired chamber (**Fig. 4D, E**). Two days later, rats were indexed again for avoidance behavior immediately following intra-CeA infusions of R121919 (0.25 μg/side) or Vehicle (Posttest 3). Non-Avoiders did not show changes in time spent in odor-paired chamber between Posttest 1 and Posttest 3 (**Fig. 4F**). On the other hand, Avoiders treated with intra-CeA R121919 (but not vehicle) showed lower avoidance behavior in Posttest 3 relative to Posttest 1 (repeated measures ANOVA, effect of Posttest, *F*_1,6_ = 22.9, *p* = 0.003) (**Fig. 4G**).

## DISCUSSION

In these experiments, we characterized behavioral and biological correlates of an avoidance phenotype in our bobcat urine stress model. We found that avoidance behavior is persistent after predator odor re-exposure and is resistant to extinction. In addition, body weight gain is lower in both Avoiders and Non-Avoiders (relative to unstressed Controls) 1 and 7 days after 1^st^ predator odor exposure, but only in Avoiders 1 day after 2^nd^ predator odor exposure. In the brain, we found that Avoiders have more c-Fos+ cells and CRF-ir than Non-Avoiders, Controls, and animals sacrificed after their 1^st^ predator odor exposure (Single Stress). We also found that high avoidance was correlated with high number of c-Fos+ cells and high CRF immunoreactivity in CeA. Finally, we showed that intra-CeA CRFR1 antagonism reverses post-stress escalation of alcohol drinking and reduces avoidance behavior in Avoiders.

The finding that both Avoiders and Non-Avoiders show decreased body weight gain relative to unstressed Controls suggests that all rats exposed to bobcat urine exhibit physiological signs of stress. Our lab previously reported that all rats exposed to predator odor exhibit increases in plasma ACTH and CORT [22]. In the current study, predator odor-exposed rats displayed blunted weight gain over the ensuing 1 and 7 days (**Fig. 1C, D**). This blunted weight gain following predator odor exposure was not accompanied by decreases in thymus weight or adrenal hypertrophy, suggesting that neuroendocrine effects of predator odor exposure may not account for the observed body weight effects. Previous work by other labs demonstrated that acute and chronic (7-14 days) predator exposure robustly activates the HPA axis and suppresses weight gain in rats but does not produce adrenal hypertrophy or thymic involution [19]. In the current study, predator odor re-exposure reduced body weight gain only in Avoiders (**Fig. 1E**), suggesting that Avoiders may be more susceptible (i.e., less resilient) to the physiological effects of repeated stress. We also report that predator odor re-exposure did not significantly alter mean group avoidance of the predator odor-paired chamber. That said, predator odor re-exposure did significantly increase avoidance of the odor-paired chamber in a subset of animals initially classified as Non-Avoiders. Previous studies have shown that repeated exposure to a predator (cat) produces HPA axis activation in rats that does not habituate [19]. This is supported by our data showing equal or exaggerated suppression of body weight gain and equal or exaggerated avoidance of odor-paired stimuli after repeated versus single stress exposure.

We report here that avoidance of predator odor-paired stimuli is resistant to extinction in Avoiders. Avoidance of the predator odor-paired chamber persisted after 3 and 6 extinction sessions, but avoidance eventually extinguished after 9 extinction sessions (**Fig. 1G**). In other animal models of PTSD, rats exposed to cats or cat collars exhibit locomotion suppression upon re-exposure to the predator-paired chamber, and this locomotor suppression has been shown to be resistant to extinction [18,23]. Humans with PTSD have extinction-resistant traumatic memories and exhibit impaired fear extinction/suppression [24]. Therefore, our predator odor model may be useful for understanding the biological factors underlying the resistance of avoidance behavior/traumatic memories to extinction.

The CeA mediates the behavioral and physiological effects of stress in rats [25]. However, whether or not acute stress exposure produces c-Fos activation in CeA seems to depend on the modality of the stressor. For instance, restraint stress, but not cat [26] or ferret odor exposure [27], increases c-Fos activation in CeA [28]. Previous studies also show that stress-induced c-Fos activation in CeA may be predictive of coping strategy. For example, rats selectively bred for high freezing and low defensive burying behaviors when presented with a shock probe (low novelty responding rats; passive/reactive copers) show more CeA c-Fos activation following footshocks than their low freezing/high defensive burying counterparts (high novelty responding rats; active copers) [29]. Here, we found that Avoiders have more c-Fos+ cells in CeA than Non-Avoiders, Controls, and Single Stress rats not indexed for avoidance, and that the number of CeA c-Fos+ cells is positively correlated with avoidance of the predator odor-paired context (**Fig. 2**), suggesting that greater CeA activation supports an Avoider phenotype/coping strategy.

CRF-CRFR1 signaling has a role in mediating stress effects on physiology and behavior. For example, systemic CRFR1 antagonism 30 min prior to or immediately after predator stress (cat exposure) blocks escalated anxiety-like behavior after stress, suggesting that CRF-CRFR1 signaling plays an important role in stress reactivity [30]. Similarly, systemic CRFR1 antagonism reduces escalated alcohol drinking following bobcat urine exposure in rats [20] or social defeat stress in mice [31], showing that CRF-CRFR1 signaling plays a role in stress-alcohol interactions. Within the brain, we predicted that the CeA is an important site in which CRF-CRFR1 signaling modulates stress reactivity and stress-induced escalation of alcohol drinking. Prior work showed that footshock stress increases CRF mRNA in CeA [32], that intra-CeA CRF receptor blockade blocks consolidation of contextual fear conditioning [33], and that acute or repeated intra-CeA CRFR1 antagonism blocks escalation of alcohol drinking in rats chronically exposed to high doses of alcohol, a procedure that usually produces affective and somatic signs of alcohol dependence [34,35]. In our lab, we previously found that intra-CeA CRF infusions produce conditioned place avoidance in stress-naïve rats and that intra-CeA CRFR1 antagonism attenuates stress-induced hyperalgesia in Avoiders [16]. Here, we report that Avoiders have greater CeA CRF-ir, CeA CRF-ir is positively correlated with avoidance behavior (**Fig. 3**), and intra-CeA CRFR1 antagonism reduces alcohol drinking and avoidance behavior in Avoiders, providing further evidence that CeA CRF-CRFR1 signaling plays a functionally relevant role in post-stress behavioral dysregulation in Avoiders.

The precise neural circuit mechanism by which CeA CRF-CRFR1 signaling contributes to avoidance behavior and escalated alcohol drinking after predator odor stress is unclear. The CeA receives CRF afferents from distal brain regions [36], but also contains local CRF neurons that release CRF within the CeA [37]. We currently do not know which CRF source contributes to the greater CeA CRF immunoreactivity in Avoiders (**Fig. 3**), and we are using circuit-based manipulations to test this question. With regard to signaling mechanisms downstream of CRF release, it is well known that CRF increases GABAergic neurotransmission in putative CeA interneurons via a CRFR1-dependent mechanism, and effect that is reproduced by alcohol bath application and is exacerbated in alcohol-dependent rats [34]. Given that intra-CeA CRFR1 antagonism blocks escalation of alcohol drinking in alcohol-dependent rats [34], mapping the circuits engaged by CeA CRFR1-dependent GABAergic transmission may be useful for understanding the mechanism underlying avoidance behavior and escalated alcohol drinking after predator odor stress.

In conclusion, the current study characterizes behavioral and biological correlates of avoidance behavior in our bobcat urine stress model and identifies CeA CRF-CRFR1 signaling as an important contributing factor to post-stress avoidance behavior and escalated alcohol drinking. Future studies will characterize the contributions of specific CRFR1-gated CeA circuits to post-stress behavioral dysregulation and extend the current findings to females.

## ACKNOWLEDGMENTS

We thank Dr. Maureen Basha for her technical assistance and guidance in dissecting adrenal and thymus glands, and Dr. Lisa Harrison-Bernard for microscope access.

